# Repurposing Niclosamide Ethanolamine for Alveolar Echinococcosis Reveals a Disconnect Between *In Vitro* Efficacy and *In Vivo* Outcome

**DOI:** 10.64898/2026.02.09.704778

**Authors:** Matías Preza, Nicole Dietrich, Pascal Zumstein, Judith Steinmann, Lea Hiller, Trix Zumkehr, Tobias Kämpfer, Marylène Chollet-Krugler, Laura Vetter, Andrew Hemphill, Sarah Dion, Britta Lundström-Stadelmann

**Affiliations:** Institute of Parasitology, Vetsuisse Faculty, University of Bern, 3012 Bern, Switzerland; Graduate School for Cellular and Biomedical Sciences, University of Bern, 3012 Bern, Switzerland; CNRS, ISCR (Institut des Sciences Chimiques de Rennes)-UMR 6226, University of Rennes, 35000 Rennes, France; Univ Rennes, Inserm, EHESP, Irset (Institut de recherche en santé, environnement et travail) - UMR_S 1085, 35000 Rennes, France; Multidisciplinary Center for Infectious Diseases, University of Bern, 3012 Bern, Switzerland

**Author notes:** Sección Biología Celular, Facultad de Ciencias, Universidad de la República, 11200 Montevideo, Uruguay (Matías Preza).

**Keywords:** Keyword: Alveolar echinococcosis, *Echinococcus multilocularis*, Niclosamide ethanolamine, Antiparasitic treatment, Drug repurposing

## Abstract

**Background:** Echinococcosis is a zoonotic disease caused by cestodes of the genus *Echinococcus*. Alveolar echinococcosis (AE), caused by *E. multilocularis*, primarily affects the liver and shows infiltrative, tumor-like growth of the metacestode stage. If untreated, AE is lethal. AE remains a neglected disease with current treatments based on albendazole or mebendazole that are parasitostatic, and not curative, underscoring the need for more effective therapies. Niclosamide is a chlorinated salicylanilide derivative with proven activities against intestinal helminths but is inactive against tissue-dwelling helminths due to poor absorption and limited bioavailability. In this study, we repurposed niclosamide ethanolamine (NEN), a formulation with improved systemic exposure, for the treatment of *E. multilocularis* infection *in vitro* and *in vivo*.

**Methodology/Principal Findings:** We assessed the *in vitro* efficacy of niclosamide and NEN against *E. multilocularis* metacestode vesicles (IC_50_<0.2 µM) and primary parasite cells (IC_50_<0.3 µM), with active concentrations largely corresponding to NEN levels reachable in the liver. Metabolic analysis suggested that NEN acts as a mitochondrial uncoupler. Electron microscopy showed that NEN-treatments induced profound structural damage in the metacestode vesicle tissue, but mitochondrial ultrastructure was not notably affected.

In mice intraperitoneally infected with *E. multilocularis*, NEN was orally administered during 9 weeks either alone, or in combination with albendazole. Pharmacokinetic analyses showed that NEN reached blood level concentrations above 1 µM. However, the parasite burden in NEN-treated mice was not significantly reduced.

**Conclusions/Significance:** Although niclosamide and NEN demonstrated potent activity against *E. multilocularis in vitro*, this efficacy did not translate in the mouse model. The lack of *in vivo* activity could be attributed to several factors such as infection model, limited drug uptake by the parasite in the animal, or the rapid metabolization of the compound. Future studies should explore novel niclosamide derivatives and formulations to enhance efficacy against AE *in vivo*.

**Author Summary:** Alveolar echinococcosis (AE) is a severe disease caused by the larval stage of the fox tapeworm *Echinococcus multilocularis*. The parasite forms tumor-like lesions in the liver and can spread to other organs. The currently licensed drugs for the treatment of AE are not always effective, require long-term use, and can cause side effects that frequently require treatment interruption. Therefore, safer and more efficacious treatment options are urgently needed.

Niclosamide is frequently applied for the treatment of adult tapeworm infections in the intestine, but its limited uptake and low biodistribution renders the compound unsuitable for systemic treatment. In this study, we tested a non-toxic salt formulation, niclosamide ethanolamine (NEN), exhibiting improved absorption. *In vitro*, NEN was highly effective against *E. multilocularis* metacestode vesicles. It induced profound structural alterations in metacestode vesicles and impaired the mitochondrial membrane potential, and thus interfering in energy production. However, NEN was not effective against AE in experimentally infected mice.

Our results suggest that NEN treatment appears promising *in vitro*, but to translate to the *in vivo* situation, new formulations and delivery strategies should be developed to increase absorption, bioavailability and metabolic stability of the compound for an effective treatment for AE.

## Introduction

Tapeworms (cestodes) are parasitic flatworms that cause significant disease in humans and domestic animals. Some, including echinococcosis, are recognized by the WHO as neglected tropical diseases (Casulli, 2020). *Echinococcus multilocularis* and *E. granulosus* are the causative agents of alveolar echinococcosis (AE) and cystic echinococcosis (CE), respectively. AE accounts for a median of 10’489 cases annually (Lundström-Stadelmann et al., 2025), whereas CE leads to at least 188’000 new cases per year, imposing 688,000 and 184,000 disability-adjusted life years (DALYs), respectively (Torgerson et al., 2015). CE is globally distributed, while AE is mainly found in continental Europe, Asia, and North America (Lundström-Stadelmann et al., 2025) and has shown a rising incidence in Europe in recent years (Casulli et al., 2025). *E. multilocularis* remains the most relevant foodborne parasite in Europe (Bouwknegt et al., 2018), and a central experimental model for *Echinococcus* and tapeworm biology.

The life cycle of *E. multilocularis* involves foxes as primary definite hosts, with other canids acting as secondary hosts. Adult worms reside in the small intestine and shed eggs into the environment via feces. Rodents serve as intermediate hosts: after ingestion, the oncosphere hatches, penetrates the intestinal wall, and reaches the liver via the bloodstream, where it develops into the metacestode, the disease-causing larval stage. Protoscoleces form within brood capsules inside the metacestode. When a definitive host consumes an infected rodent, protoscoleces reach the intestine and develop into adult worms, completing the life cycle. In intermediate hosts, but also aberrant hosts like humans, dogs or non-human primates, metacestodes proliferate infiltratively in the liver, undergo tumor-like and expansive growth, and form lesions that finally cause AE (Kern et al., 2017). Metacestodes are characterized by a vesicular structure. The outer layer of these vesicles is formed by the laminated layer (LL), a carbohydrate-rich entity that defines the host-parasite interface. Attached to the inner lining of the LL is the tegument, a syncytial layer with microvilli-like protrusions forming the connection to the LL. The germinal layer (GL) further inwards is a multicellular parasite tissue, and it is comprised of numerous cell types, including muscle cells, nerve cells, glycogen storage cells, and stem cells. Metacestodes are filled with vesicle fluid, which is secreted by GL cells (Brehm and Koziol, 2017).

Treatment options against AE are limited. Curative therapy relies on the complete surgical removal of the parasite, which is often impossible due to the localization and extent of the lesion, as well as the need for access to appropriate surgical infrastructure. Surgery is accompanied by benzimidazole drug therapy, and in cases where radical surgery cannot be performed, patients rely solely on long-term drug treatment with benzimidazoles, typically albendazole (ABZ) or mebendazole. These drugs are parasitostatic and fail to eliminate stem cells, so the parasite persists and the disease is not cured. Longterm, in many cases life-long, drug treatment is needed, often accompanied by adverse effects (Grüner et al., 2017; Horton, 1997). Therapeutic drug-level monitoring is necessary to balance efficacy and toxicity, but this is not consistently available in low-resource settings (Lundström-Stadelmann et al., 2025). Given these limitations, there is an urgent need for new and more effective and safer drug therapies for AE.

Different strategies exist to identify new therapeutic options against AE. The tumor-like proliferative capacity of *E. multilocularis* enables large-scale in *vitro* metacestode vesicle cultivation. Based on this, an *in vitro* drug screening cascade was established (Lundström-Stadelmann et al., 2019). It allows to assess compound activities against whole metacestode vesicles and GL cell cultures including stem cells and studies on the mode of action of active compounds. Promising candidates are subsequently evaluated for drug efficacy in established *in vivo* mouse models of AE (Lundström-Stadelmann et al., 2019; Wang et al., 2024).

Several compounds have shown *in vitro* activity against *E. multilocularis*, including nitazoxanide (Stettler et al., 2003), amphotericin B (Reuter et al., 2003b), buparvaquone (Rufener et al., 2018), metformin (Loos et al., 2020), atovaquone (Enkai et al., 2020), 2-Deoxy-D-glucose (Xin et al., 2022), kinase inhibitors (Koike et al., 2022), endochin-like quinolones (Chaudhry et al., 2022), and mefloquine (Lundström-Stadelmann et al., 2020; Memedovski et al., 2023), among others. Few of these repurposed compounds were also tested in the *in vivo* AE mouse model, but none of them exhibited *in vivo* efficacy that was comparable to ABZ. To date, the polyene macrolide antibiotic Amphotericin B has remained the sole compound with reported benefit in some cases of human AE as salvage treatment in patients that could not be treated with benzimidazoles (Jelicic et al., 2023; Reuter et al., 2003a). Thus, the discovery of more effective and improved therapeutic options against AE is still urgently needed.

Niclosamide is an FDA-approved anthelmintic (Chen et al., 2018), but its poor gastrointestinal absorption and unfavorable pharmacokinetics and biodistribution has prevented effective targeting of tissue-dwelling parasites. To overcome this limitation, we have focused here on a formulation of niclosamide with an improved systemic bioavailability: niclosamide ethanolamine (NEN). NEN is repurposed from the field of oncology research (Chen et al., 2017; Jiang et al., 2022; Wang et al., 2022) and reaches high liver concentrations of 1.5 µM within 4 h in mice treated with 40 mg/kg (Tao et al., 2014a). In this study, we demonstrate the profound activity of NEN against *E. multilocularis* metacestodes *in vitro*, show that NEN affects the mitochondrion as a primary target, and investigate the pharmacokinetics and *in vivo* efficacy of NEN in the AE mouse model.

## Materials and Methods

### Chemicals, tissue culture reagents, and cells

All chemicals were purchased from Sigma-Aldrich (Buchs, Switzerland) and all plastic ware was obtained from Sarstedt (Sevelen, Switzerland), if not stated otherwise. Dulbeccos’s modified Eagle medium (DMEM) and penicillin and streptomycin (10’000 Units/mL penicillin, 10’000 μg/mL streptomycin) were purchased from Gibco (Fisher Scientific AG, Reinach, Switzerland). NEN was obtained from 2A Biotech (Lisle, IL, USA). Fetal bovine serum (FBS) and Trypsin/EDTA (0.05% Trypsin/0.02% EDTA) were purchased from Bioswisstec (Schaffhausen, Switzerland). Murine Hepa 1-6 cells were kindly provided by Magali Roques (Institute of Cell Biology, University of Bern). Reuber rat hepatoma (RH) cells (H-4-II-E) were purchased from ATCC (Molsheim Cedex, France).

### Parasite maintenance and *in vitro* culture of metacestode vesicles

*E. multilocularis* metacestodes (isolate H95) were maintained by serial intraperitoneal passage in BALB/c mice. Metacestodes proliferated *in vivo* for 3-4 months, after which animals were euthanized, and the parasite tissue was resected. *In vitro* cultures were set up and maintained as previously described (Kaethner et al., 2023). Briefly, the resected parasite material was pressed through a conventional mesh strainer and incubated overnight at 4 °C in PBS supplemented with 100 U/mL penicillin, 100 µg/mL streptomycin, 20 µg/mL levofloxacine and 10 µg/mL tetracycline. Afterwards, the metacestode vesicles were co-cultured with RH cells at 37°C in DMEM, 10% FBS, 100 U/mL penicillin, 100 μg/mL streptomycin, and 5 μg/mL tetracycline. Until further use of metacestode vesicles in experiments, culture media was refreshed weekly, and freshly detached RH cells were added to the cultures.

### Assessment of NEN-induced metacestode vesicle damage by phosphoglucose isomerase assay

Damage of *E. multilocularis* metacestode vesicles was tested by the damage marker release assay based on phosphoglucose isomerase (PGI) measurements, as previously described (Stadelmann et al., 2010). Briefly, after 10-12 weeks of *in vitro* culture, metacestode vesicles were washed extensively in PBS, and resuspended in DMEM without phenol red, with penicillin (100 U/mL) and streptomycin (100 µg/mL). The metacestode vesicle suspension in medium was distributed into 48 well plates (1 ml/well). Serial 1:3 dilutions (10 µM to 13.7 nM) of niclosamide, NEN, and ethanolamine as a control, were tested in technical triplicates, all in 0.1% DMSO. Negative controls received only DMSO (0.1%). 0.1% Triton X-100 was used as a positive control inducing total metacestode vesicle damage. Drug-treated metacestode vesicles were incubated at 37 °C in a humidified atmosphere with 5% CO_2_. Images of each well were acquired on days 0 and 5 using a Nikon SMZ18 stereo microscope (Nikon, Basel, Switzerland) at 7.5 × magnification. PGI release was evaluated from supernatant samples taken at day 5 of drug-incubation and activity was measured indirectly as previously described (Stadelmann et al., 2010). Half maximal inhibitory concentration (IC_50_) values were calculated from four biologically independent experiments using RStudio (Ritler et al., 2023) by the statistical model Weibull type 2 (3 parameters).

### E. multilocularis GL cell isolation

The isolation of GL cells was performed as previously described (Kaethner et al., 2024). Conditioned medium (cDMEM) was obtained by the incubation of 10^6^ RH cells in 50 mL medium for six days and 10^7^ RH cells in 50 mL medium for four days, both of them in DMEM, 10% FBS, 100 U/mL penicillin, 100 μg/mL streptomycin, and 5 μg/mL tetracycline, at 37°C under a humid atmosphere with 5% of CO_2_ as described.

Six-months-old metacestodes vesicles from *in vitro* cultures were washed extensively with PBS, followed by immersion in distilled water to deplete them from remanent RH cells, and washed again with PBS. The PBS was removed, and metacestode vesicles were broken by passing them through a 1 mL pipette, and the suspension was centrifuged at 600 × g for 6 min at RT to separate the tissue from the vesicle fluid. Vesicle fluid was removed, and the tissue washed extensively with PBS. The tissue was treated with trypsin 0.05%/EDTA 0.02% for 30 min at 37°C, followed by repeated cycles of shaking and filtering using a 30 µm mesh (Sefar AG, Heiden, Switzerland). The filtered material was collected and centrifuged 50 × g for 30 sec at RT to remove calcareous corpuscles. Afterwards the cells were concentrated with a centrifugation of 600 × g for 10 min at RT. The pellet was resuspended in cDMEM. The OD_600_ of the resulting cell suspension was measured three times using a 1:100 dilution, and the average value was calculated. An OD_600_ of 0.1 was defined as 1 arbitrary unit (AU) per µL of the undiluted cell suspension, corresponding to 0.97 ± 0.11 µg of total protein for GL cells, as determined by a bicinchoninic acid (BCA) protein assay (Zumstein et al., 2025)). 1,000 AU were incubated in 5 mL of cDMEM overnight under nitrogen atmosphere at 37 °C. The next day the cells were resuspended and re-incubated under nitrogen atmosphere at 37 °C for 3 h before use for experiments.

### Viability assays on *E. multilocularis* GL cells

For drug efficacy testing, GL cells were centrifuged and resuspended in cDMEM. 15 AU were distributed to each well of a 384 well plate (Huberlab, Aesch, Switzerland). Niclosamide, NEN, and ethanolamine were added to final concentrations ranging from 10 µM to 4.9 nM using 1:2 serial dilutions, in technical quadruplicates, and incubated in a microaerobic chamber (85% N_2_, 10% CO_2_, 5% O_2_), humid atmosphere. Images of each well were acquired on day 5, using a Nikon TE2000E microscope connected to a Hamatsu ORCA ER camera at 40 × magnification. For cell viability evaluation, the ATP content in each well was measured by CellTiter-Glo (Promega, Dübendorf, Switzerland) including 1% Triton X-100. The luminescent emission was measured on a HIDEX Sense microplate reader (Hidex, Turku, Finland). The viability was calculated relative to the signal of the DMSO 0.1% controls. IC_50_ values were calculated from four biologically independent experiments in RStudio (Ritler et al., 2023), applying the 5-Parameter Logistic Model.

### Toxicity of niclosamide and NEN on mammalian cells

This assay was adapted from the protocol described by Rufener et al. (2018). RH cells and Hepa 1-6 cells were used for resazurin-based assessment of mammalian cell toxicity, both at confluent and pre-confluent (proliferative) stages. Cells were seeded in 96-well plates at densities of 50’000 cells/well (confluent) or 5’000 cells/well (proliferating), respectively, in DMEM supplemented with 10% FBS, 100 U/mL penicillin, 100 μg/mL streptomycin, and 5 μg/mL tetracycline. Cells were incubated overnight at 37°C, 5% CO_2_ for confluent conditions or for 5 h for non-confluent conditions, before addition of drugs. NEN, niclosamide, and ethanolamine were added in serial 1:3 dilutions (from 120 µM to 54.9 nM). After 5 days of drug incubation cell viability was assessed by removing the culture medium, washing with PBS, and adding resazurin at 10 µg/mL in PBS. Fluorescence was measured at 595 nm using a Hidex multilabel reader at 22°C, time 0 and 50 min. Four independent experiments were done for each cell line, and each experiment contained three technical replicas. For each cell line, dose-response curves were fitted with RStudio (Ritler et al., 2023) applying the 4 Parameter Weibull type 2 Model. Selectivity Index was calculated as follows: IC_50_ (host cells)/ IC_50_ (*E. multilocularis* GL cells).

### Transmission electron microscopy of drug-treated metacestode vesicles

Metacestode vesicles were isolated and cultured under the same conditions as described in the section above for PGI assay. Metacestode vesicles were treated with 300 nM NEN for 12, 24, 48 h, and 5 days, while negative controls received DMSO at 0.1%. Following drug treatments, metacestode vesicles were broken by passing them through a 1 mL pipette. They were washed 3 times with PBS, followed by one wash in cacodylate buffer (100 mM, pH 7.3), and samples were fixed overnight at 4 °C in cacodylate buffer containing 2% glutaraldehyde. Subsequent post fixation was done at RT for 2 h in cacodylate buffer containing 2% osmium tetroxide at RT. The specimens were then washed with water, and stepwise dehydrated using a graded series of ethanol solutions (30-50-70-90-100%). Finally, the samples were embedded in EPON-812 epoxy resin for 48 h at RT, and the resin was polymerized by an overnight incubation at 60 °C. Ultrathin sections were cut on a ultramicrotome (Reicher & Jung, Vienna, Austria) and were loaded onto 200 mesh nickel grids (Plano GmbH, Marburg, Germany). They were then contrasted using Uranyless™ and lead citrate solutions (Electron microscopy Sciences, Hatfield, PA, USA), and specimens were imaged on a FEI Morgagni transmission electron microscope (TEM) equipped with a Morada digital camera system (12 Megapixel) operating at 80 kV.

### Seahorse assays to detect NEN-induced effects on oxygen consumption in GL cells

Mitochondrial respiration of *E. multilocularis* GL cells was assessed using the Seahorse Mito Stress Test, following the manufacturer’s protocol for the Seahorse XFp Extracellular Flux Analyzer (Agilent Technologies, Basel, Switzerland), with slight modifications as described previously (Zumstein et al., 2025). As GL cells are not adherent, we coated Seahorse XFp cell culture miniplates with Cell-Tak (22.4 µg/mL, Fisher Scientific, Schwerte, Germany) and distributed 100 AU of GL cells per well, in Seahorse DMEM supplemented with 1 mM pyruvate, 2 mM glutamine, and 10 mM glucose. For measurement of respiration, we injected oligomycin to a final concentration of 1.5 µM in the first port. With the injection of the second port, we tested one of the following three conditions: Carbonyl cyanide 4-(trifluoromethoxy)phenylhydrazone (FCCP, final concentration 0.5 µM), NEN (final concentration 1 µM) or DMSO (final concentration, 0.1%) as a negative solvent control. We performed experiments with three independent isolations of GL cells. In each experiment two cartridges were tested, in each cartridge we included two technical replicates for each condition (thus a total of four replica each). The relative oxygen consumption rate (OCR) increase was calculated by subtracting the OCR measured before the addition of FCCP, NEN, or DMSO from the OCR measured after the addition, and then normalizing the result by division with the pre-injection OCR value × 100. For statistical analysis we performed ANOVA followed by post-hoc test using Tukey’s honest significant difference test to identify pairwise differences between the incubation conditions of three independent biological replica in RStudio.

### Tetramethylrhodamine ethyl ester assay to detect NEN-induced effects on mitochondrial potential in GL cells

To assess effects of NEN treatment on the mitochondrial potential of GL cells, tetramethylrhodamine ethyl ester (TMRE) assay was employed. The TMRE dye accumulates in mitochondria in a membrane potential–dependent manner, allowing assessment of mitochondrial membrane potential (Scaduto and Grotyohann, 1999). GL cells were cultured for at least one week to allow aggregate formation (Zumstein et al., 2025). Aggregates were then centrifuged at 100 × g for 3 min at RT and resuspended in pre-warmed cDMEM (see above). 100 AU in 180 µL were seeded per well in round-bottom 96-well plates. Drug stocks in DMSO were diluted 1:100 in cDMEM to prepare 10X working solutions; 20 µL per well yielded a final volume of 200 µL. Cells were incubated for 1 hour at 37°C under nitrogen atmosphere. TMRE (10 mM) was diluted in cDMEM to 2 µM, added to a final concentration of 0.1 µM per well, and incubated for 30 min at 37°C under nitrogen atmosphere. A negative control without TMRE was included. Cells were gently resuspended, transferred to 1.5 mL centrifuge tubes, washed three times with 1 mL PBS (centrifuged at 300 × g for 1 min per wash), and finally resuspended in 50 µL PBS. The entire volume was transferred to a 384-well plate, centrifuged at 300 × g for 3 min, and analyzed on a Nikon Eclipse Ti 2 Spinning Disk (Cicero) microscope. Fluorescence in aggregates was analyzed using Fiji (ImageJ, Version 2.14.0). Statistical analysis was performed as for the Seahorse assay using three biologically independent experiments.

### Ethical statement and mouse maintenance

*In vivo* studies were conducted in accordance with the Swiss animal protection law (TschV, SR 455). The study received approval from the Animal Welfare Committee of the Canton of Bern (licence numbers BE12/24 and BE2/22). Female and male BALB/c mice, 8 weeks old, were obtained from Charles River Laboratories (Sulzheim, Germany). Before the experiments, the animals were acclimatized for 2 weeks. These mice were used for drug efficacy testing when they reached 11 weeks of age and weighed about 20 grams. The mice were kept in BlueLine Cage Typ II (Tecniplast), with up to five mice per cage. The cages included nestlets (Plexx, Elst, Netherlands), a tunnel (Zoonlab, Castrop-Rauxel, Germany), and a mouse house (Tecniplast, Gams, Switzerland) as enrichment. They were maintained under a 12-hour light/dark cycle, with a controlled temperature of 21°C–23°C, and a relative humidity of 45%–55%. Food and water were provided *ad libitum*.

### Assessment of NEN in a secondary infection model of AE in mice

24 female and 12 male 8-weeks old BALB/c mice were infected by intraperitoneal injection of 100 µL of *E. multilocularis* material from *in vitro* cultures as previously described (Karpstein et al., submitted). For this, metacestode vesicles maintained in culture for over 6 months were washed three times with PBS. Following removal of the PBS, vesicles were broken by passing them through a 20G syringe. The vesicle fluid and the parasite tissue were separated by centrifugation (300 × g, 3 min, at RT) and the volume of the pellet was topped up with the same amount of PBS to generate a final suspension for infection. Mice were randomized into new cages after one week (3 mice per cage, with females and males housed separately). Randomization was performed using an Excel spreadsheet with the “RAND” function. Treatment was initiated at four weeks post-infection. Four experimental groups were used; all treated twice per day with 6 h between treatments. Each group consisted of six females and three males (thus n=9 per treatment arm): Vehicle (only vehicle twice per day), ABZ (first dose ABZ 200 mg/kg, second dose vehicle), NEN (NEN 40 mg/kg twice per day), and combination treatment group (first dose ABZ 200 mg/kg plus NEN 40 mg/kg, second dose NEN 40 mg/kg). All drug treatments were performed by micropipette-guided drug administration (MDA) (Scarborough et al., 2020). To prepare the vehicle and respective drug-emulsions, 3 volumes of condensed milk (MIGROS, Kondensmilch, gezuckert, Migros, Zurich, Switzerland) were diluted in 10 volumes of autoclaved tap water. The milk solution was mixed with the drugs using a glass bead of 5 mm, and a Homogeniser, FastPrep®-24 (MP Biomedicals, Europe) with the default settings (20 sec 4 m/s). Animal compliance to MDA method was evaluated each day by assigning a score from 0 to 6 to every mouse, reflecting its performance over time (see S1 Table).

Mouse body weight was measured weekly to adjust dosages according to the weight. During the experiment, drug-induced side effects, including alterations in fur appearance, grooming behavior, posture, and overall activity levels, were carefully assessed and scored using a standardized score sheet. After 9 weeks of treatment, all mice were euthanized by CO_2_ inhalation. The parasite material was dissected and weighted in a petri dish. Pearson correlation analysis was performed in RStudio to assess the relationship between parasite weight and blood NEN concentration (see below).

### Blood sampling and serum sample preparation for HPLC

In weeks 3 and 7 of treatment, blood samples were taken from the tail vein at 1, 2 and 4 h after the first daily dose. Different mice were used as representatives for each time point within each treatment group. For each group and time point, samples were obtained from two females and one male. At euthanasia, an additional blood sampling was performed, which was taken from the heart, again 1, 2 and 4 h after treatment and from different animals (two females and one male per time point per treatment group). The blood was left at RT for 30 min, followed by centrifugation at 3000 × g at 4°C. Serum was then collected from the supernatant, and samples stored at −80°C until NEN concentrations were measured as described below.

### UV-HPLC analysis of serum samples

To prepare serum samples for HPLC analysis, 60 to 80 μL of serum were mixed with 2 μL methanol, 100 µL acetonitrile and 10 μL naproxen internal standard (1500 ng/mL in acetonitrile, IS). The mixture was vortexed for 1 min and centrifuged at 12,000 × g for 20 min at RT. Calibration standards were prepared by spiking blank serum (30 µL) with NEN at 250, 500,1000, 1250 and 1500 ng/mL and processed identically. After centrifugation, 10 µL aliquots of supernatant were stored at −80°C until UV-HPLC analysis. For HPLC-DAD analysis, 10 µL of each serum extract and standard were injected into an HPLC-DAD system (Shimadzu®, Marne La Vallée, France, LC-20 AD SP, injector SIL-20A HI, column oven CTO-20A, diode array detector SPD-M20A) equipped with an InfinityLab Poroshell SB-C18 column (150 x 4.6 mm, i.d. 4 µm, Agilent Technologies^TM^). Mobile phase A:H_2_0 + 0.1 % formic acid; mobile phase B: acetonitrile + 0.1 % formic acid. Gradient: 50% B (0–2 min), 50–70% B (2–5 min), 70% B (5–6 min), 70–50% B (6–6.5 min), 50% B (6.5–9 min). Flow rate: 1 mL/min; column temperature 40 °C. Analysis was done using the Lab Advisor software (Agilent Technologies^TM^, Dako, Les Ulis, France). Peak detection was carried out online using a diode array detector. Naproxen (internal standard) was detected at 220 nm (retention time ∼3.9 min). NEN was detected at 335 nm (retention time ∼7.7 min). Identification in samples was based on retention time matching against standards. Quantification used the ratio of NEN peak area to internal standard peak area and the calibration curve: NEN area/ISaArea = 0,0006 C_NEN_ – 0,2407 (R^2^ = 0,944; C_NEN_ in ng/mL).

### Manuscript preparation

The graphs were made in RStudio using the packages *ggplot2*, the final figures for the manuscript were designed in Adobe Illustrator 29.8.2.

The linguistics of selected sentences was improved using ChatGPT (version 5.2).

## Results

### Niclosamide and NEN damage *E. multilocularis* metacestode vesicles and impair the viability of isolated GL cells *in vitro*

We applied the PGI assay, a damage marker release assay, to assess the efficacy of niclosamide and NEN on whole *E. multilocularis* metacestode vesicles *in vitro*. After five days of drug treatment, niclosamide and NEN, but not ethanolamine alone, had induced significant amounts of PGI release into the medium supernatant, indicative for physical damage of metacestode vesicles (Figure 1A). This was also seen by light microscopy, showing morphological changes in drug-treated metacestode vesicles (Figure 1B). Clear drug-induced morphological alterations were visible at 370 nM and higher, consistent with the IC_50_ values of 166.3 ± 77.9 nM for niclosamide and 168.3 ± 81.0 nM for NEN (Table 1). Niclosamide and NEN also strongly impaired ATP production and thus the viability of GL cells at slightly higher concentrations (Figure 1C) with IC_50_ values of 257.4 ± 51.0 nM for niclosamide, and 230.1 ± 54.8 nM for NEN (Table 1). Microscopy confirmed this effect, with typically formed GL aggregates present in DMSO and ethanolamine-treated cultures, but not in GL cultures exposed to niclosamide and NEN from 625 nM and higher (Figure 1D). Ethanolamine alone neither induce any PGI release nor affect GL cell viability.

**Fig 1.**
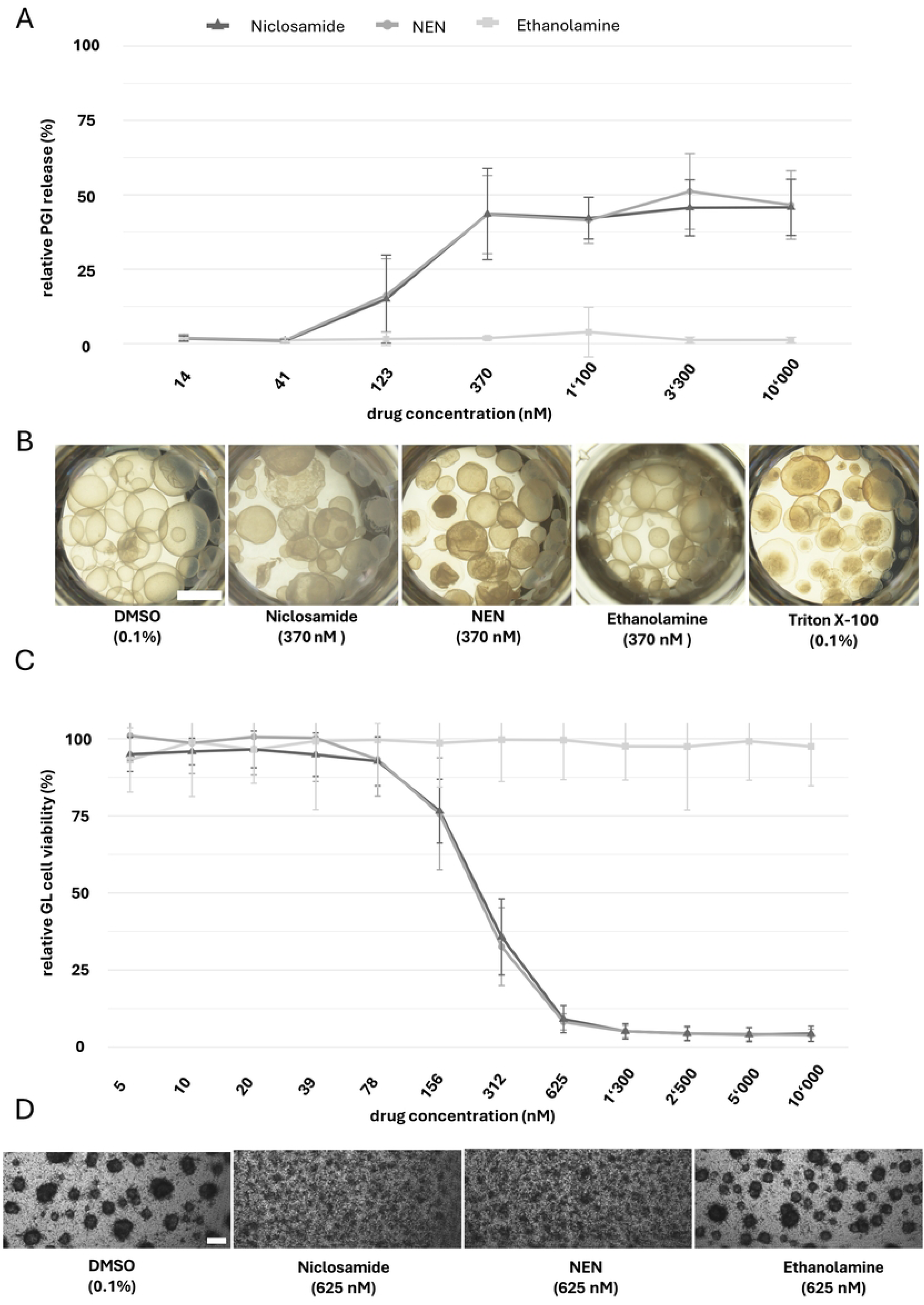
Niclosamide and NEN damage *E. multilocularis* metacestode vesicles and reduce the viability of GL cells *in vitro*. A, PGI release assay was employed after 5 days of metacestode vesicle treatment to assess the efficacy of niclosamide (dark grey) and NEN (grey) - both at concentrations ranging from 10’000 to 13.7 nM. DMSO solvent controls are subtracted, and all data is presented relative to the PGI values obtained in the presence of 0.1% Tx-100. Four independent assays were conducted each with technical triplicates. Mean values and SDs from all experiments performed are shown. B, representative light microscopic images of metacestode vesicles treated with 370 nM niclosamide, NEN, and ethanolamine, DMSO (0.1%) and Tx-100 (0.1%). Scale bar = 2 mm. C, GL cell viability as assessed by CellTiter Glo assay upon treatment of GL cells for 5 days with a concentration range (10’000 to 4.9 nM) of niclosamide (dark grey), NEN (grey) or ethanolamine (light grey). Data is shown relative to values obtained upon treatment with 0.1% DMSO (100% viability), and mean values and SDs of four independent experiments are shown, all conducted in technical quadruplicates. D, Representative photomicrographs of drug-treated and control-treated GL cells are shown. Scale bar = 100 µm.

**Table 1.** Summary of IC_50_ values of niclosamide and NEN against *E. multilocularis* and mammalian cells. IC_50_ values were calculated for PGI damage-marker release assay on whole metacestode vesicles. Viability was calculated for isolated *E. multilocularis* GL cells, RH cells, and Hepa 1-6 cells. Mean values ± SDs of four independent experiments are shown. Ethanolamine alone did not induce any damage in the metacestode vesicles nor effects on viability. ND: not determined.

| Assay and cell type | Niclosamide |  | NEN |  |  |
| --- | --- | --- | --- | --- | --- |
|  | Mean IC<sub>50</sub> values ± SD (nM) | Selectivity Index comparison to GL cells | Mean IC<sub>50</sub> values ± SD (nM) | Selectivity Index comparison to GL cells | *p*-values niclosamide vs NEN |
| PGI assay *E. multilocularis* metacestode vesicles | 166.3 ± 77.9 | - | 168.3 ± 81.0 | - | 0.97 |
| Viability of isolated GL cells of *E. multilocularis* | 257.4 ± 51.0 | - | 230.1 ± 54.8 | - | 0.49 |
| Confluent RH cells | ND | ND | ND | ND | ND |
| Proliferating RH cells | 2'173.8 ± 511.7 | 8.45 | 2'278.3 ± 335.2 | 9.90 | 0.75 |
| Confluent Hepa 1-6 cells | 3'311.1 ± 2'042.3 | 12.86 | 6'165.8 ± 1'177.4 | 26.80 | 0.06 |
| Proliferating Hepa 1-6 cells | 1'306.2 ± 326.6 | 5.07 | 851.6 ± 386.2 | 3.70 | 0.12 |

### Niclosamide and NEN show higher selectivity for *E. multilocularis* than for mammalian cells *in vitro*

Mammalian cell toxicity was assessed using confluent and subconfluent/proliferating rat hepatoma (RH) and murine hepatoma (Hepa 1-6) cell cultures. IC_50_ values could not be calculated for confluent RH cells, as up to a concentration of 120 µM, no complete loss of viability was detected. For proliferating RH cells, the IC_50_ values were 2’173.8 ± 511.7 nM for niclosamide and 2’278.3 ± 335.2 nM for NEN (Table 1). Hepa 1-6 cells were overall more sensitive to both compounds. On confluent Hepa 1-6 cells, the IC_50_ values were 3’311.1 ± 2’042.3 nM for niclosamide and 6’165.8 ± 1’177.4 nM for NEN. On proliferating Hepa 1-6 cells, the IC_50_ values were 1’306.2 ± 326.6 nM for niclosamide, and 851.6 ± 386.2 nM for NEN (Table 1). In summary, Table 1 highlights that *E. multilocularis* metacestode vesicles and GL cells are more sensitive to niclosamide and NEN than mammalian cells (selectivity indexes in the range of 3.7 to 26.8). In proliferating RH cells, selectivity indices were 8.45 for niclosamide and 9.90 for NEN. In Hepa 1-6 cells, higher selectivity was observed in confluent cultures, with selectivity indices of 12.86 for niclosamide and 26.80 for NEN, compared to proliferating cultures (5.07 and 3.70, respectively). Niclosamide and NEN were similarly active in all assays, thus, the formulation did not lead to any loss of drug activity. Therefore, subsequent analyses focused on NEN.

### NEN induces ultrastructural changes in *E. multilocularis* metacestode vesicles as observed by transmission electron microscopy

Treatment of metacestode vesicles with 0.1% DMSO does not affect the structural integrity of the GL, tegument, and LL, and microtriches were abundantly present at the GL-LL boundary (Figure 2A-C). After 12, 24, and 48 h of treatment with 300 nM NEN, metacestode vesicles remained structurally unaltered (S1 and S2 Figures). Notable alterations became evident only after 5 days of NEN treatment, indicating that the drug requires a prolonged period of continuous exposure to exert its effects. The observed changes seen in the GL included the accumulation of lipid droplets and black inclusions, increased numbers of vacuoles, some with content of unknown nature, and a partial deterioration of the GL tissue integrity (Figure 3A-C). While the tegument remained largely unaltered, microtriches were partially shortened, and the matrix of the LL contained an increased number of extracellular vesicle-like structures. However, the undifferentiated cells (or stem cells), characterized by a large nucleus and numerous mitochondria, remained morphologically largely unaltered by these treatments (Figure 3D). The presence of mitochondria with an intact electron-dense matrix and clearly discernible cristae indicated that the viability of undifferentiated cells was not affected by the NEN treatment (Figure 3E).

**Fig. 2.**
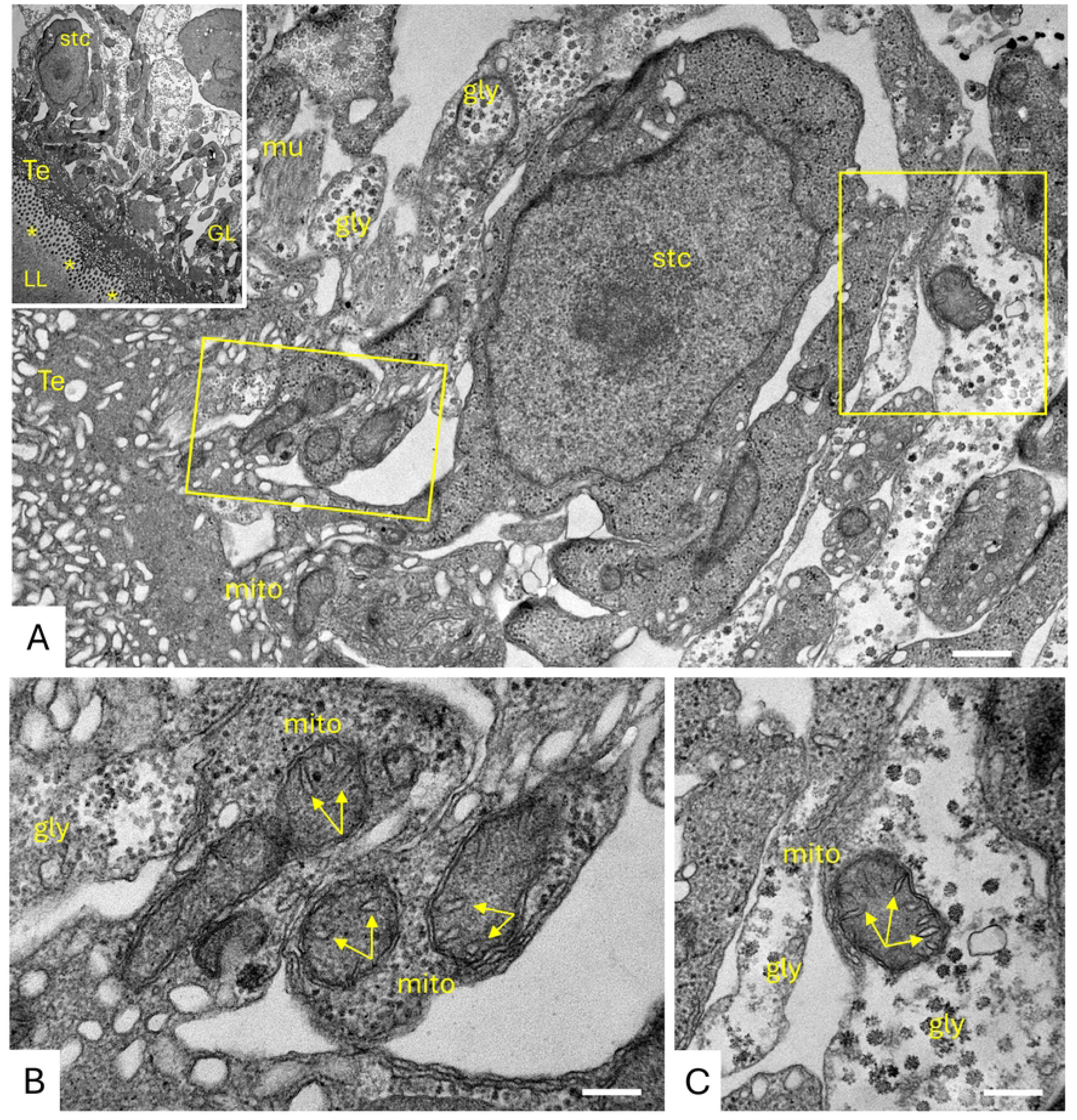
TEM of *in vitro* cultured *E. multilocularis* metacestode vesicles without drug treatment. A, B, and C show control metacestode vesicle tissue treated after 5 days of culture in 0.1% DMSO. The insert in A shows a lower magnification view with laminated layer (LL), tegument (Te) and germinal layer (GL). Microtriches cross-sections are depicted with asterisks (*). A subtegumentary cyton (stc) is shown at higher magnification in A; nuc = nucleus, gly = glycogen storage cells, mito = mitochondrion. The boxed areas in A are shown at higher magnification in C and D, with the mitochondrial cristae indicted by arrows. Scale bars: A = 1.2 µm; B = 0.5 µm; C = 0.8 µm

**Fig 3.**
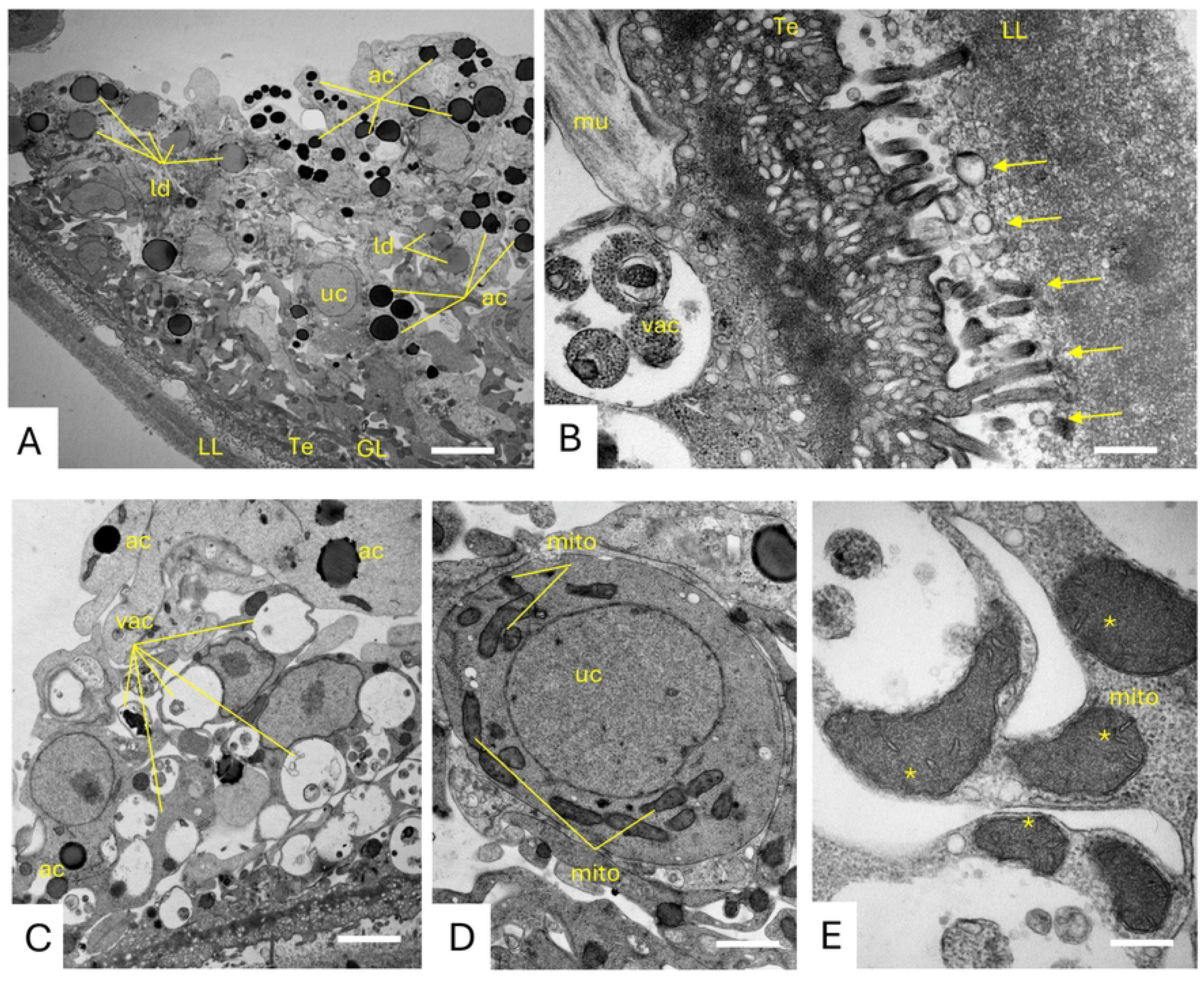
TEM of *E. multilocularis* metacestode vesicles treated with NEN. Metacestode tissue treated with 300 nM NEN during 5 days. Treatments results in the accumulation of lipid droplets (ld) and black inclusions, most likely acidocalcisomes (ac) (A), increased numbers of cytoplasmic vacuoles (B and C), some deterioration of the microtriches (arrows) and increased numbers of small vesicles being secreted into the LL (B). However, undifferentiated cells shown in D), and respective mitochondria (E) remained largely unaltered. Scale bars: A = 4.8 µm; B = 0.7 µm; C = 5 µm; D = 1.4 µm; 0.3 µm. GL = germinal layer; LL = laminated layer; Te = tegument; uc = undifferentiated cell; ga = Golgi apparatus; mito = mitochondrion; gsc = glycogen storage cells; mu = muscle cells; arrows = microtriches; * = cristae; ld = lipid droplets, ac = acidocalcisomes; * = cristae; arrows: microtriches plus extracellular-like vesicles; vac = vacuolization.

### NEN treatment of *E. multilocularis* GL cells affects the mitochondrial membrane potential

The effects of NEN were evaluated on isolated *E. multilocularis* GL cells using Seahorse and TMRE assays. Seahorse analysis showed a significant increase in OCR after NEN treatment (*p*-value = 1.0E-06), comparable to the standard uncoupler FCCP (*p-*value = 1.7E-03), while DMSO had no effect (Figure 4A, S2 Table). TMRE staining was strong in control cells, indicative for a marked mitochondrial membrane potential, but reduced after NEN exposure (*p-*value = 2.3E-04), with an even stronger decrease than that induced by FCCP (*p*-value = 1.6E-04, Figure 4B, C). Overall, these results indicate that NEN acts as an uncoupler and affects mitochondrial function in parasite cells.

**Fig 4.**
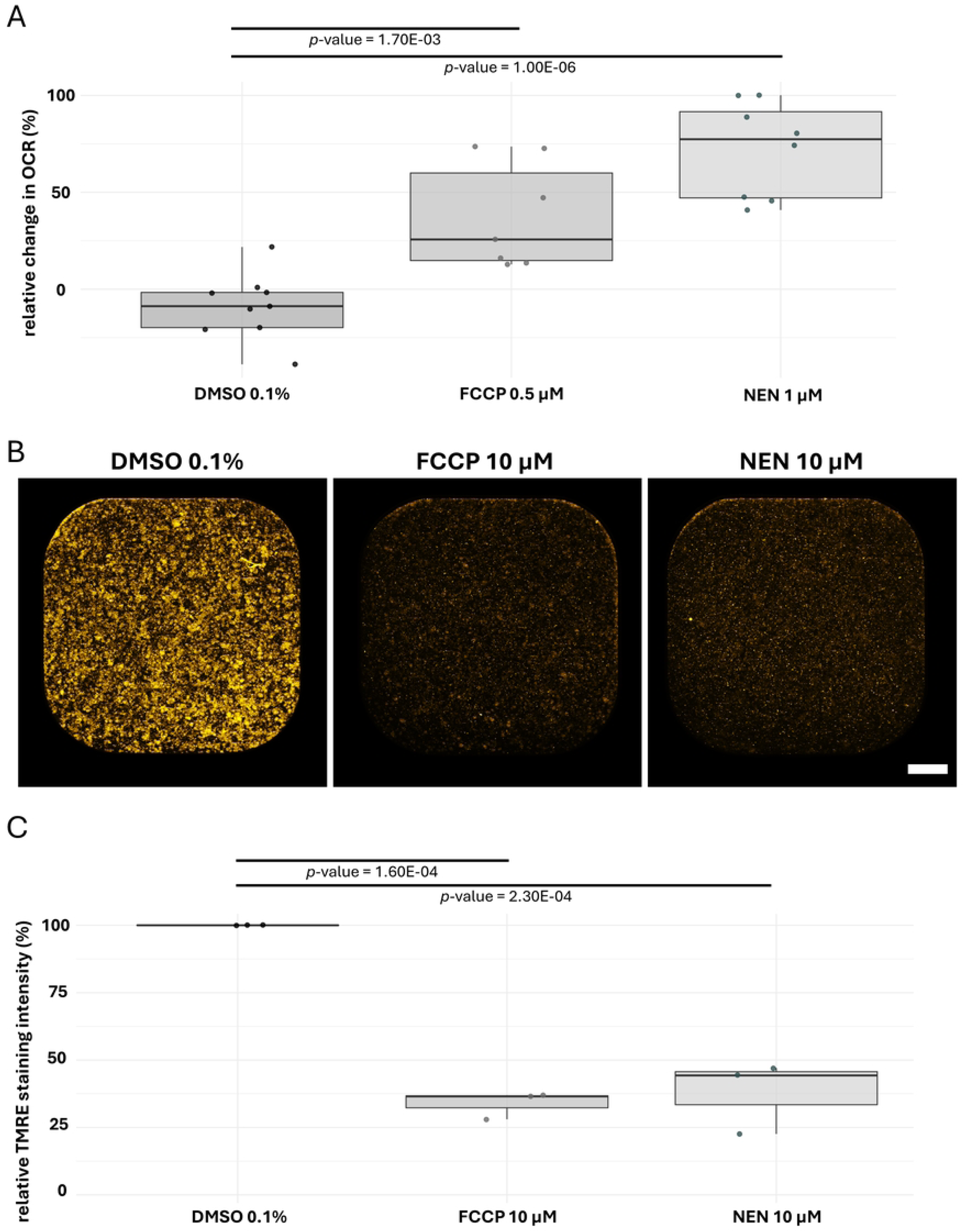
NEN acts as a mitochondrial uncoupler in *E. multilocularis* GL cells. A, Relative change in the oxygen consumption rate (OCR) upon treatment of GL cells with DMSO (0.1%, negative control), FCCP (0.5 µM, positive control), or NEN (1 µM). Data points from three independent experiments with four technical replica each are shown, including corresponding box plots. B, Representative pictures of GL cells treated with DMSO (0.1%), FCCP (10 µM), or NEN (10 µM) stained with TMRE. Scale bar = 500 µm. C, Relative TMRE staining intensity of GL cells treated with DMSO (0.1%, negative control), FCCP or NEN (both 10 µM). Group comparisons were performed using one-way ANOVA followed by Tukey’s post-hoc multiple comparisons test; adjusted *p*-values are shown. Data points from three independent experiments are shown with corresponding box plots.

### NEN is not effective in a murine intraperitoneal AE infection model, even though niclosamide blood levels are high

To assess the *in vivo* efficacy and translational potential, treatments with NEN and NEN combined with ABZ were tested in an intraperitoneal AE infection model in mice. These treatments were comparatively assessed to standard ABZ treatment.

Based on its previously reported pharmacodynamic properties (Tao et al., 2014b), NEN was administered twice daily. To reduce stress for both animals and researchers, drug administration was performed by MDA. The success of MDA was monitored daily by scoring the performance of each mouse over time (S1 Table). Most mice readily consumed the full dose while being held only by the tail, and accepted the treatment completely voluntarily, without any handling, often actively searching for the dose as soon as the cage was opened. The overall median score for voluntary drug consumption by MDA was 5. For the vehicle and ABZ groups the median score was 6. In the NEN and ABZ + NEN treated groups, male mice were less willing to take the drug voluntarily, and for this reason, these individuals were treated by conventional oral gavage. These groups had a median score of 5. No adverse side effects were observed in any of the animals during the study. Two mice were euthanized two weeks prior to the planned endpoint due to rapid parasite progression, reaching termination criteria (one in the vehicle group and one in the NEN group).

The parasite burden in all treatment groups was assessed after 9 weeks of treatment. Compared to the vehicle-treated group (4.87 ± 3.37 g), NEN treatment did not significantly reduce the growth of the parasite tissue, and a tendency toward a higher parasite mass was observed, although this difference was not statistically significant (6.31 ± 4.34 g, *p-*value = 7.71E-01). No differences between mouse sexes were observed, and no significant treatment effects were detected when analyzed separately by sex (S3 Table). ABZ treatment significantly reduced the parasite burden to 1.16 ± 1.33 g (*p*-value = 2.12E-02). A similar reduction was observed in the group receiving a combination of NEN and ABZ 1.01 ± 0.92 g (*p*-value = 1.57E-02). However, no synergistic effect was noted, as the combination therapy did not outperform ABZ alone (*p*-value = 0.99, Figure 5, S3 Table).

**Fig 5.**
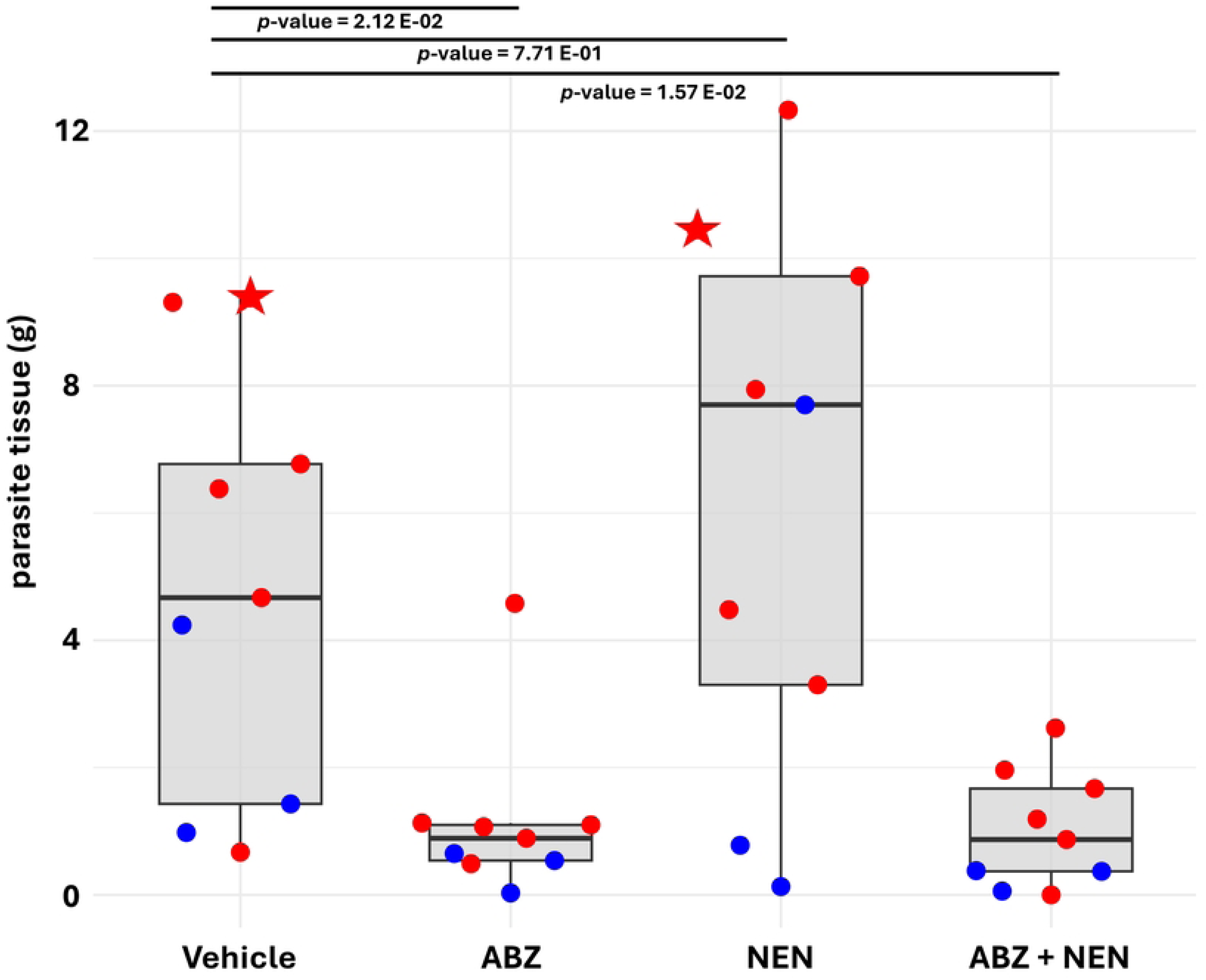
Efficacy of NEN on AE in mice. A, Intraperitoneally *E. multilocularis* infected mice were treated orally with either NEN (40 mg/kg twice per day), ABZ (200 mg/kg once per day), a combination of both (ABZ + NEN), or the vehicle control (condensed milk). Each treatment arm consisted of 9 animals (6 females, in red, and 3 males, in blue) and respective parasite weights (g) after 9 weeks of treatment are shown for each individual with corresponding box plots for each treatment group. Group comparisons were performed using the Kruskal–Wallis test followed by Dunn’s post-hoc multiple comparisons test with Benjamini–Hochberg correction; adjusted *p*-values are shown. Two mice were euthanized before the endpoint of the experiment and they are depicted with asterisks.

To assess systemic exposure to NEN, we measured its serum concentration in mice treated with NEN alone or in combination with ABZ at 1, 2 and 4 h post treatment. Serum levels were analyzed on treatment weeks 3, 7 and 10. We detected NEN concentrations higher than 1 µM at every time point analyzed in both groups, thus well above the IC_50_ values determined by PGI assay *in vitro* (S4 Table). However, variations in the results were also seen, with samples under the limit of detection for each time point. Similar values were obtained in samples obtained from uninfected mice after the first and second treatment of NEN or NEN and ABZ (S4 Table). In general, lower average NEN concentrations were detected at 4 h post-treatment, suggesting rapid systemic depletion. Using Pearson correlation analysis, NEN treatment alone showed a weak negative correlation between NEN concentration and parasite tissue mass (r = –0.226), whereas the ABZ+NEN combination produced a substantially stronger negative correlation (r = –0.652), suggesting enhanced efficacy when both drugs were co-administered.

## Discussion

Alveolar echinococcosis is a neglected disease causing over ten thousand new human cases each year worldwide (Lundström-Stadelmann et al., 2025). Despite the severity of the infection, fully effective pharmacological treatment remains unavailable. Here, we evaluated the efficacy of niclosamide and its ethanolamine-salt formulation NEN against AE. Both niclosamide and NEN showed strong activity against *E. multilocularis* metacestode vesicles and isolated GL cells *in vitro*. Damage-marker release assays yielded IC_50_ values of ∼ 150 nM, and GL cell viability assays produced IC_50_ values of ∼250 nM. These concentrations indicate a greater potency than several previously reported *in vitro*-active compounds, including amphotericin B (tested at 2.71 μM, (Reuter et al., 2003b)), nitazoxanide (tested at 3.25-32.5 µM (Stettler et al., 2003)), metformin (tested at 10 mM, (Loos et al., 2020)), and mefloquine (with IC_50_ >30 µM, (Lundström-Stadelmann et al., 2020)). However, as different experimental approaches were used across studies, direct comparisons of effective concentrations should be interpreted with caution. The *in vitro* activity of niclosamide and NEN falls closer to that of buparvaquone (IC_50_: 2.87 µM in PGI assay, Rufener et al., 2018)), and ELQ-400 (IC_50_: 1.06 µM in PGI assay, Chaudhry et al., 2022)), both targeting mitochondrial function via *bc*_1_ complex inhibition.

Our findings indicate that the antiparasitic effect of NEN is, at least partially, driven by mitochondrial uncoupling. This uncoupling effect of niclosamide has been already documented in multiple biological systems (Williamson and Metcalf, 1967; Tao et al., 2014b). Related salicylanilides, including rafoxanide, closantel and oxyclozanide, show flukicidal activity (Mohammed-Ali and Bogan, 1987), and disrupt oxidative phosphorylation in trematodes (Fairweather and Boray, 1999). The recent identification of the uncoupler ESI-09 as a compound with profound activity against *E. multilocularis* (Zumstein et al., 2025), along with other reports of uncouplers, further supports mitochondrial uncoupling as a promising avenue for therapeutic development against parasitic flatworms. This also aligns with the reduced ATP levels we observed in NEN-treated GL cells and supports previous findings pointing to mitochondrial metabolism as a promising therapeutic target in *E. multilocularis* (Enkai et al., 2020). Unexpectedly, TEM analysis did not reveal structural alterations in parasite mitochondria, consistent with previous observations for the uncoupler ESI-09. Notably, treatments with both NEN and ESI-09, and several other compounds active against metacestodes, led to the accumulation of extracellular-like vesicles in the matrix of the LL (Zumstein et al., 2025). This response may represent a broader stress reaction of the parasite and warrants further investigation.

Compounds structurally related to niclosamide have also been evaluated against *E. multilocularis*. Both nitazoxanide and MMV665807, showed potent *in vitro* activity against metacestode vesicles and GL cells (Stadelmann et al., 2016; Stettler et al., 2003). However, despite this promising *in vitro* efficacy, *in vivo* studies revealed limited therapeutic effects (Stadelmann et al., 2016; Stettler et al., 2004). These findings highlight that compounds with high potency *in vitro* must also exhibit favorable pharmacokinetic, metabolic stability and biodistribution properties *in vivo*, and the use of delivery strategies that enhance these properties could render them effectiv*e*. The mouse model for AE treatment remains challenging to use. *E. multilocularis* establishes slowly, and treatments require long-term repeated dosing, which is stressful for both animals and researchers. To reduce this burden, we used a semi-voluntary drug administration method via MDA. This refinement is in line with the 3R principles for animal experimentation, which are gaining increasing importance in experimental research (Lauwereyns et al., 2024). Overall, the method worked well, as reflected by the mostly cooperative behavior of the mice and the clear treatment response to ABZ.

Despite the excellent efficacy of NEN *in vitro*, the parasite burden was not reduced upon *in vivo* treatment. The most plausible explanation is poor drug penetration into parasite tissues within the peritoneal cavity. Measured serum concentration at 1, 2 h post-treatment was generally above the levels required for *in vitro* activity. The most plausible explanation is that serum levels did not translate into effective exposure at the actual infection site. However, we were not able to implement measurement of drug level in parasite tissue. The serum levels obtained in this study are consistent with previously reported niclosamide concentrations in mice (Tao et al., 2014b). In the infection model used here, parasites develop within the peritoneal cavity, where local drug exposure is likely lower than in well-perfused tissues supplied by peripheral blood vessels. This limitation underscores the importance of infection models that more closely reflect the natural hepatic localization of *E. multilocularis* and allow improved assessment of drug distribution to parasite lesions. An organotypic infection model based on oral egg-infection, or a mouse model applying direct inoculation of metacestodes into the liver *in vivo*, allowing the parasite to grow in hepatic tissue (Romig and Bilger, 1999; Siles-Lucas and Hemphill, 2002) would better reflect the natural infection site and might allow higher drug accumulation at the lesion, potentially improving therapeutic performance. However, access to *E. multilocularis* eggs is limited, and the associated biosafety risks are substantial. Thus, despite its constraints, the intraperitoneal infection model represents the advanced, disseminated stage of disease and remains a standard experimental system (Albani et al., 2015; Liu et al., 2021; Rufener et al., 2018; Wang et al., 2016).

Niclosamide is currently being investigated as a therapeutic agent across several disease areas, including multiple cancer types (Chae et al., 2017; Jiang et al., 2022; Shangguan et al., 2020), COVID (Singh et al., 2022), and diverse infections and inflammatory conditions (Tao et al., 2014a; Esmail et al., 2021; Z. Liu et al., 2024), but its development for clinical use is limited by poor systemic exposure and low metabolic stability (Choi et al., 2021). Addressing these limitations will require optimized formulations or structural analogues that improve solubility, stability, and tissue distribution. Several efforts move in that direction, such as the acyl analogue DK-520 (Mook et al., 2015). Additional strategies such as nanoparticle-based delivery systems (Lohiya and Katti, 2021), liposomal formulations (Y. Liu et al., 2024), or prodrugs (Kim et al., 2023) aim to increase drug levels at target tissues with the goal to improve therapeutic performance. New compounds should be evaluated in combination with current standard treatments ABZ or mebendazole, which remain the most effective options for AE. Combination therapies may enhance antiparasitic efficacy through additive or synergistic effects while enabling dose reduction, shortening of treatment durations, and thus lowering toxicity. In clinical settings, ABZ would typically be continued rather than replaced, increasing the translational relevance of such strategies. Future work should also assess safety, particularly hepatic tolerance, given the liver tropism of both parasite and drugs involved.

## Conclusion

Overall, niclosamide and its ethanolamine formulation NEN show clear antiparasitic activity *in vitro*, yet in the *in vivo* mouse model efficacy against *E. multilocularis* metacestodes remains limited, a challenge also seen with several other compounds tested previously. AE continues to be a neglected disease, and the lack of commercial incentive continues to slow therapeutic innovation. Nevertheless, the broader therapeutic interest in niclosamide is likely to accelerate the development of improved formulations, analogues, and prodrugs that could also benefit AE treatment. Assessing nanoparticle-based delivery is a particularly relevant direction and prior work on mefloquine-loaded PLGA-PEG-COOH nanoparticles (Autier et al., 2024) illustrates the feasibility of encapsulating poorly soluble antiparasitic compounds to enhance pharmacokinetics and tissue targeting. Addressing poor bioavailability, rapid systemic clearance, and limited tissue penetration will be essential to bridge the gap between *in vitro* potency and *in vivo* efficacy. Advances in drug delivery and pharmacokinetic optimization remain central to realizing the potential of niclosamide and related molecules within future treatment strategies for AE.

## Acknowledgements

The authors thank the animal caretakers from the Vetsuisse Faculty of the University of Bern; their professionalism and kindness ensured that this work was carried out under optimal conditions. The authors also thank Magali Roques (Institute of Cell Biology, University of Bern) for kindly sharing the Hepa 1-6 cells.

## Funding statement

This study was supported by grants to BLS from the Uniscientia Foundation and the Gottfried and Julia Bangerter-Rhyner Foundation, as well as the Swiss National Science Foundation (SNSF) grant number 320030-227438 and 320030-236056.

The funders had no role in study design, data collection, analysis and interpretation, decision to publish, or preparation of the manuscript.

## Author contributions

Matías Preza: Conceptualization, Data Curation, Formal Analysis, Investigation, Methodology, Supervision, Validation, Visualization, Writing – Original Draft Preparation, Writing – Review & Editing.

Nicole Dietrich: Investigation.

Pascal Zumstein: Investigation, Methodology, Validation, Writing – Review & Editing. Judith Steinmann: Investigation.

Lea Hiller: Investigation.

Trix Zumkehr: Investigation, Methodology, Writing – Review & Editing. Tobias Kämpfer: Investigation, Methodology, Writing – Review & Editing.

Marylène Chollet-Krugler: Investigation, Methodology, Writing – Review & Editing. Laura Vetter: Investigation.

Andrew Hemphill: Data Curation, Formal Analysis, Investigation, Visualization, Writing – Review & Editing.

Sarah Dion: Investigation, Methodology, Writing – Review & Editing

Britta Lundström-Stadelmann: Conceptualization, Formal Analysis, Funding Acquisition, Investigation, Project Administration, Resources, Supervision, Visualization, Writing – Original Draft Preparation, Writing – Review & Editing.

---

**S1 Fig. TEM of *E. multilocularis* metacestode vesicles treated with 300 nM NEN for 12 h.** After 12 h of treatment with NEN 300 nM *in vitro* no alterations are observed in metacestode vesicle structure. B shows a zoom view of the zone marked in A with an unalerted mitochondria. In B, details from the zone marked in A with an arrowhead are shown where the microtriches are shown in detail. E, zoom in of the cells marked in D where normal undifferentiated cells are shown. CGL = germinal layer; LL = laminated layer; Te = tegument; uc = undifferentiated cell; mito = mitochondrion; gsc = glycogen storage cells; mu = muscle cells; arrows = microtriches; arrows: microtriches plus secreted vesicles. Scale bars: A = 2 µm; B = 0.5 µm; C = 0.4 µm; D = 2.8 µm; E = 1 µm.

**S2 Fig. TEM of *E. multilocularis* metacestode vesicles treated with 300 nM NEN for 24 and 48 h.** After 24 (A and B) and 48 (C - F) h of treatment with NEN 300 nM *in vitro* no alterations are observed in metacestode vesicle structure. B, zoom view from the microtriches structure marked in A with an arrowhead. D, zoom view from the region marked in C. F, zoom view from the region marked in E. GL = germinal layer; LL = laminated layer; Te = tegument; uc = undifferentiated cell; ga = Golgi apparatus; mito = mitochondrion; gsc = glycogen storage cells; arrows = microtriches; * = cristae; ld = lipid droplets, arrows: microtriches plus secreted vesicles. Scale bars: A = 1 µm; B = 0.5 µm; C = 2.2 µm; D = 0.7 µm; E = 2.8 µm.; F = 0.9 µm.

**S1 Table. MDA scoring of each mouse during morning and afternoon treatments.** For each individual treatment every mouse was scored from 0 to 6 depending on their will to take the treatment.

**S2 Table. Seahorse XFp Mito Stress Test results.** OCR values over time of the Seahorse assays using GL cells of *E. multilocularis*.

**S3 Table. *In vivo* results.** Parasite material (g) recovered for each mouse of the different groups in the endpoint of the *in vivo* experiment.

**S4 Table - Concentration (µM) of NEN in mouse serum.** Mice were treated with NEN (NEN 40 mg/kg) or with ABZ+NEN (ABZ 200 mg/kg plus NEN 40 mg/kg), blood samples were taken 1, 2 and 4 h after treatment.

## Notes

### Competing Interest Statement

The authors have declared no competing interest.

